# omicsGAT: Graph Attention Network for Cancer Subtype Analyses

**DOI:** 10.1101/2022.06.08.495390

**Authors:** Sudipto Baul, Khandakar Tanvir Ahmed, Joseph Filipek, Wei Zhang

**Affiliations:** Department of Computer Science, Genomics and Bioinformatics Cluster, University of Central Florida, Orlando, FL 32816, USA

**Keywords:** Graph Attention Network, Single-cell RNA-seq, Patient Stratification, Cancer Outcome Prediction

## Abstract

**Motivation:** The use of high-throughput omics technologies is becoming increasingly popular in all facets of biomedical science. The mRNA sequencing (RNA-seq) method reports quantitative measures of more than tens of thousands of biological features. It provides a more comprehensive molecular perspective of studied cancer mechanisms compared to traditional approaches. Graph-based learning models have been proposed to learn important hidden representations from gene expression data and network structure to improve cancer outcome prediction, patient stratification, and cell clustering. However, these graph-based methods cannot rank the importance of the different neighbors for a particular sample in the downstream cancer subtype analyses. In this study, we introduce omicsGAT, a graph attention network (GAT) model to integrate graph-based learning with an attention mechanism for RNA-seq data analysis. The multi-head attention mechanism in omicsGAT can more effectively secure information of a particular sample by assigning different attention coefficients to its neighbors.

**Results:** Comprehensive experiments on The Cancer Genome Atlas (TCGA) breast cancer and bladder cancer bulk RNA-seq data, and primary diffuse gliomas single-cell RNA-seq data validate that (1) the proposed model can effectively integrate neighborhood information of a sample and learn an embedding vector to improve disease phenotype prediction, cancer patient stratification, and cell clustering of the sample. (2) The attention matrix generated from the multi-head attention coefficients provides more useful information compared to the sample correlation-based adjacency matrix. From the results, we can conclude that some neighbors play a more important role than others in cancer subtype analyses of a particular sample based on the attention coefficient.

**Availability and implementation:** Source code is available at: https://github.com/CompbioLabUCF/omicsGAT

**Supplementary information:** Supplementary data are available at *BioRxiv* online.

## Introduction

Cancer is a complex and heterogeneous disease with hundreds of types and subtypes spanning across different organs, tissues and have origins in various cell types (1, 2). For example, breast cancer is highly heterogeneous with different subtypes that lead to varying clinical outcomes including prognosis, response to treatment, and changes of recurrence and metastasis (3–5). Hence, cancer subtype prediction and cancer patient stratification have been the subject of interest to clinicians and patients for many decades. Powered by the high-throughput genomic technologies, the mRNA sequencing (RNA-seq) method is capable of measuring transcriptome-wide mRNA expressions and molecular activities in cancer cells (6, 7). Bulk RNA-seq data provides a view of the average gene expression level of an entire tissue sample instead of differentiating among cell types within the sample. Whereas, single-cell RNA-seq (scRNA-seq) provides opportunities to explore gene expression profiles at the single-cell level. These will enable predicting the changes of expression level at a large scale so as to better understand the biological mechanism that leads to cancer.

The high-throughput RNA-seq datasets show quantitative measures of more than tens of thousands of mRNA isoforms for a cohort of hundreds or thousands of samples (e.g., patients, cells). However, due to the unavoidable sample heterogeneity or experimental noise in the data, extracting biological valuable information and discovering the underlying patterns from the data is becoming a serious challenge to computational biologists (8). While hundreds of computational methods have been developed for cancer subtype prediction/identification (9, 10) and patient stratification (11) using RNA-seq data (12), network analysis of sample similarities has largely been ignored in most methods. Graph-based neural network (GNN) and network-based embedding models recently have shown remarkable success in learning network topological structures from large-scale biological data (13–15). On another note, the self-attention mechanism has been extensively used in different applications including bioinformatics (16–18). This mechanism allows inputs to interact with each other and permits the model to utilize the most relevant parts of the inputs to improve the performance of the deep learning models. The self-attention mechanism was combined with the graph-structured data by Veličković et al. (19) in Graph Attention Networks (GAT). This GAT model calculates the representation of each node in the network by attending to its neighbors, and it uses the multi-head attention to further increase the representation capability of the model (20). It applies varied attentions to the neighbors; therefore, find the most important neighbors of a sample rather than giving all of them the same importance. This model has been successfully applied on various tasks including text classification (21), node classification (22), social influence analysis (23), recommendation system (24), etc. The GAT model has also been applied to bioinformatics applications including drug-target interaction prediction (25), drug-microbe interaction prediction (26), gene essentiality prediction (27), etc.

Inspired by the GAT for capturing node dependencies in a wide range of domains, we proposed omicsGAT model and applied it on cancer samples with RNA-seq data. First, we introduced the model in Methods section. Next, we tested omicsGAT on The Cancer Genome Atlas (TCGA) breast invasive carcinoma (BRCA) data collections (28) and urothelial bladder carcinoma (BLCA) data collections (29) for cancer subtype prediction and cancer patient stratification, respectively (Section F). Then, omicsGAT was applied on 2,458 cells from six primary diffuse gliomas with K27M histone mutations (H3K27M) for cell clustering (Section G). Last, we discussed and interpreted the results based on the sample-by-sample attention matrix generated from the omicsGAT model in the Discussion section.

## Methods

In this section, we first introduce our proposed framework, omicsGAT, which generates embeddings from gene expression data to be used in downstream classification and clustering. We extended the GAT model (19) to better fit our tasks of disease outcome prediction and subtype stratification. Then, we discuss the baseline models used to compare and validate the performance of omicsGAT followed by the details of evaluation metrics used in this study.

### A. Graph Attention Network

The omicsGAT model architecture builds on the concept of the self-attention mechanism. In omicsGAT, embedding is generated from the gene expression data assuming that the samples (i.e., patients or cells) with similar features (gene expressions) are expected to have similar disease outcomes or cell types, and connected to each other. Hence, network information is injected into the model using the adjacency matrix to take these connections into consideration. However, all connected neighbors of a target sample should not get equal attention in generating the embedding for that sample. A particular neighbor of a target sample can contribute more to its subsequent prediction or clustering which cannot be accurately apprehended by similarity metrics. Therefore, to capture the importance of each neighbor on a sample, the omicsGAT model automatically assigns different attentions to the neighbors of that sample for singular head while generating the embedding. Moreover, to consider the impact of different types of information secured from the neighbors and stabilize the learning process, the above procedure is repeated multiple times in parallel employing several heads (independent attention mechanisms) in a multi-head framework.

The mathematical notations used to explain omicsGAT are summarized in Table 1. Let *n* be the number of samples (e.g., patients, cells) and *m* be the number of features (e.g., genes) representing each sample. The input feature matrix is given by ***X*** = [***x***_1_, ***x***_2_, …, ***x***_*n*_], where ***x*** ∈ ℝ^1×*m*^ represents a sample vector. ***A*** be the *n* × *n* adjacency matrix (includes self-connections) built based on the pairwise correlation between the samples. Suppose, the set of neighbors for a sample ***x***_*i*_ is denoted by 𝒩_*i*_. Depending on the number of neighbors |𝒩_*i*_ |to be kept for a sample, the connections with high correlation scores are kept (assigned a value of 1) and the others are discarded (assigned a value of 0). The adjacency matrix is binarized as it will be used to mask the attention coefficients in later part of the model. Self-connections are applied to integrate the information from the samples themselves in their embeddings. While generating the embedding of sample ***x***_*i*_, the attention given to it from its neighbor ***x***_*j*_ for a single head can be calculated as

**Table 1.** Mathematical notations for omicsGAT

| Name | Definition |
| --- | --- |
| $n$ | number of samples (i.e., patients or cells) |
| $m$ | number of features (i.e., genes) |
| $p$ | embedding size for a single head |
| $h$ | number of heads |
| $\mathbf{X} \in \mathbb{R}^{n \times m}$ | input feature matrix |
| $\mathbf{A} \in \mathbb{R}^{n \times n}$ | correlation-based adjacency matrix of samples |
| $\mathbf{W} \in \mathbb{R}^{m \times p}$ | weight matrix of a single head |
| $\mathbf{a} \in \mathbb{R}^{2p \times 1}$ | attention weight matrix of a single head |
| $\boldsymbol{\alpha} \in \mathbb{R}^{n \times n}$ | attention coefficients of a single head |
| $\mathbf{Z} \in \mathbb{R}^{n \times ph}$ | embedding matrix learned from the model |

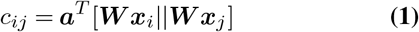

where ***W*** ∈ ℝ^*p*×*m*^ and ***a*** ∈ ℝ^2*p*×1^ are learnable weight parameters of a single head which are shared across all the samples and *p* is the embedding size. ∥ and. ^*T*^ symbols denote the concatenation and transposition operations of the matrices respectively. The calculated attention values are passed through a *LeakyReLU* activation function. Then the structural information of the network is introduced by masking the attention values using the adjacency matrix. Only the attention values where a connection is present between the nodes (samples) in the adjacency matrix ***A*** are kept and all the remaining values are made zero. After that, the attention coefficient for a neighbor ***x***_*j*_ is calculated using *Softmax* function which follows the equation below:

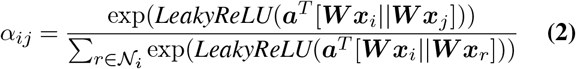

The attention coefficients calculated for all of the neighbors of ***x***_*i*_ using equation (2) are leveraged to calculate its final embedding for a single head

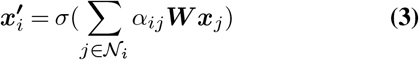

where *σ* is a non-linear activation function. Note that the sample ***x***_*i*_ is also included in its neighbors since self-connections are used in the adjacency matrix.

In a multi-head attention network, each head has a separate attention mechanism with its own weight matrix ***W*** and attention vector ***a***. Outputs generated by all the heads for one particular sample are concatenated to generate the final embedding vector of that sample. This is done to stabilize the learning process while generating the embedding. It is similar to the mechanism used by Vaswani et al. (16) in self-attention. Hence, the output embedding from the first part of our model for ***x***_*i*_ is given by:

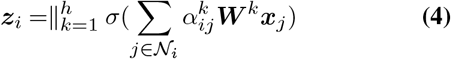

where *h* is the number of heads. The output projected in the embedding space is represented by ***Z*** ∈ ℝ^*n*×*ph*^ and embedding for one sample is ***z*** ∈ ℝ^1×*ph*^. The generated embeddings are then used in separate models for classification and clustering tasks. The overall framework of our proposed pipeline is illustrated in Figure 1.

**Fig. 1.**
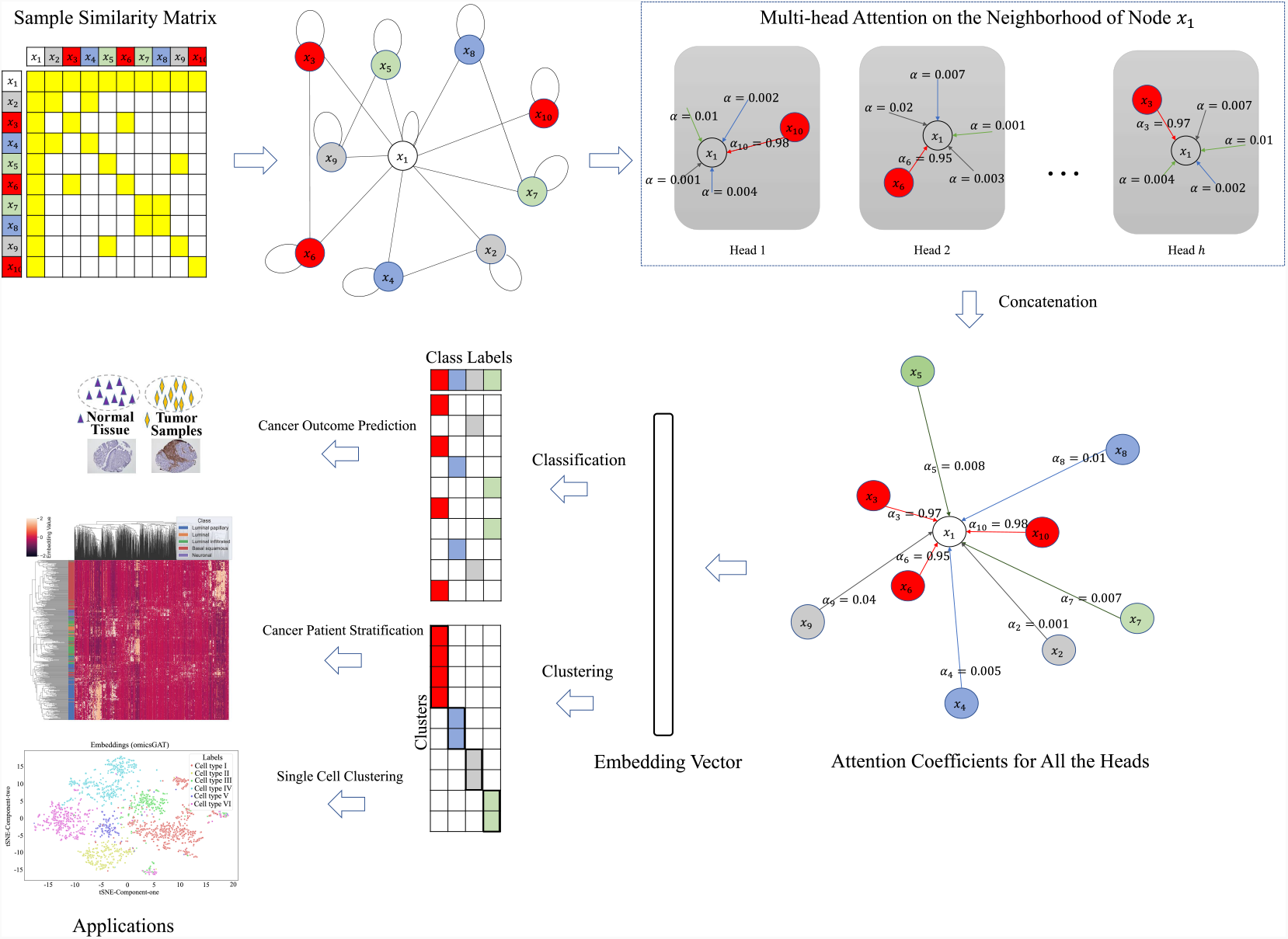
Workflow of omicsGAT. For a sample ***x***_1_, based on the input feature matrix and adjacency matrix, each head calculates the attention given to ***x***_1_ from its neighbors separately. The embeddings produced by all heads are concatenated together to generate the final embedding for ***x***_1_ which is then used for classification or clustering of ***x***_1_.

### B. omicsGAT Classifier

omicsGAT Classifier is a unified model that passes the embedding ***Z*** generated from the first part of our pipeline described in Section A through three fully connected (FC) layers. Let the number of classes for the classification task be *c*. The first two layers converts ***Z*** ∈ ℝ^*n*×*ph*^ to 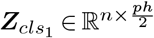 and then to 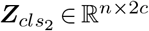 respectively. The output layer transforms 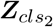 into ***Y***_*cls*_ ∈ ℝ^*n*^×^*c*^, where 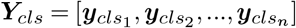 represent the classification outcomes. Each layer can be formulated as

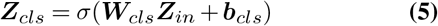

where ***Z***_*cls*_ and ***Z***_*in*_ are the output and input matrices, ***W***_*cls*_ is the learnable weight, and ***b***_*cls*_ is the bias vector of a particular layer. *σ* denotes the activation function which is *ReLU* for the first two layers and *Softmax* for the output layer.

Let the ground truth labels for *n* samples be 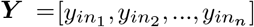. In order to calculate the overall loss of the model, Negative Log Likelihood (NLL) loss function is applied, formulated as follows:

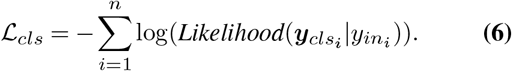

*ℒ*_*cls*_ is minimized to train the unified omicsGAT Classifier framework.

### C. omicsGAT Clustering

For clustering, we propose a two-step omicsGAT Clustering framework. The first step is an autoencoder that generates the gene expression embedding in an unsupervised approach, and the second step is a hierarchical clustering model. omicsGAT described in Section A serves as the encoder in the autoencoder architecture whereas a four layers fully connected neural network is constructed as the decoder. The output ***Z*** ∈ ℝ^*n*×*ph*^ from the omicsGAT encoder is fed into the first layer of the decoder. The output of the consecutive FC layers are 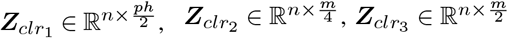, and ***Y***_*clr*_ ∈ ℝ^*n*×*m*^ respectively. Each layer can be formulated as

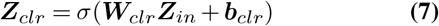

where ***Z***_*clr*_ and ***Z***_*in*_ are the output and input matrices respectively, ***W***_*clr*_ is the learnable weight, and ***b***_*clr*_ is the bias vector of a particular layer of the decoder. For the first three layers, *σ* denotes the activation function *ReLU*, and no activation function is used in the final layer.

The output, projected back to the input feature space by the decoder, is given by 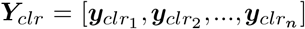. Mean squared error (MSE) is employed to calculate the reconstruction loss as follows:

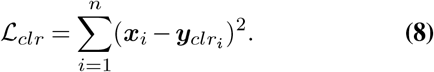

*ℒ*_*clr*_ is minimized to train the autoencoder and an embedding is generated as output from the trained encoder. The embedding is then fed into the second step of omicsGAT Clustering, a hierarchical clustering model implemented using the *scikit-learn* package (30). It stratifies the input samples into the defined number of clusters by assigning each sample to a group based on the similarity of the generated embedding of that sample with that of the other samples in the group.

### D. Baseline Models used for Comparison

#### D.1. Baselines for Classification Tasks

Support Vector Machine (SVM), Random Forest (RF), Deep Neural Network (DNN), and Graph Convolutional Network (GCN) are used as baselines to evaluate and compare the performance of omicsGAT Classifier. The baselines are built using several Python open-source library packages including *Scikit-learn* (30) and *Pytorch* (31).

SVM and RF are two of the most widely used machine learning models. In this study, ‘rbf’ kernel is applied for SVM. Hyperparameters for RF, including the number of trees, split criterion, maximum depth of the tree, maximum number of features considered for split, are also tuned. The Deep Neural Network model consists of three fully connected linear layers with first two of them followed by the *ReLU* activation function. For better evaluation of our model by comparing it to a similar graph-based deep learning model, we follow the Graph Convolution Network (GCN) proposed by Kipf and Welling (32). The GCN model is composed of four graph convolution layers. The correlation-based adjacency matrix ***A*** is used as neighborhood information in the GCN model. The hyperparameters for all of these models were tuned on the validation set using grid search.

#### D.2. Baselines for Clustering Tasks

To evaluate the embedding learned from omicsGAT, we use the clustering results of raw features and their PCA components as baselines. Hierarchical and k-means clusterings are employed for the baselines, i.e., components learned using PCA or the raw features are fed into the clustering models as input. Moreover, for a better interpretation of our model, the attention coefficients from each of the heads are extracted to build up the attention matrix which will be described in the Discussion section. The correlation-based adjacency matrix ***A*** is used as baseline to evaluate the attention matrix. Hierarchical clustering is applied on the attention matrix and adjacency matrix, both of which represent the relation among the samples.

### E. Evaluation Metrices

In this section, we define three evaluation metrics used in this study implemented using the *scikit-learn* library of Python. The Area Under the Receiver Operating Characteristic Curve (AUC) is used for comparison of the classification models. It is defined as the area under the curve plotted using True Positive Rate (*precision*) along the y-axis and False Positive Rate (1-*specificity*) along the x-axis. The Normalized Mutual Information (NMI) and Adjusted Rand Index (ARI) are applied to evaluate the clustering methods both of which have a range from 0 to 1, where 1 means perfect clustering and 0 means totally random.

## Experiments

We carried out experiments on TCGA RNA-seq datasets and H3K27M gliomas scRNA-seq data to evaluate the performance of omicsGAT in this section. In the first part, we performed experiments with omicsGAT for cancer outcome prediction on TCGA breast cancer dataset and cancer patient stratification on TCGA bladder cancer dataset (Section F). In the later part, omicsGAT was applied on scRNA-seq data for single cell clustering analysis (Section G).

### F. Experiments on TCGA Cancer Patient Samples

#### F.1. Datasets and Preprocessing

The proposed framework, omicsGAT, was tested on TCGA breast invasive carcinoma (BRCA) (28) and urothelial bladder carcinoma (BLCA) (29) datasets. The RNA-seq mRNA expression dataset of each cancer type was downloaded from UCSC Xena Hub (33). *log*2(*x* + 1) transformed mRNA expression was used in the analyses. The clinical information of the two cancer studies was downloaded from cBioPortal (34). The BRCA dataset consists of 411 patient samples and 20,351 genes for each sample. Similarly, the BLCA dataset consists of 426 patient samples and 20,531 genes for each sample.

#### F.2. omicsGAT Improved Overall Cancer Outcome Prediction

We designed two tasks on TCGA BRCA mRNA expression data to evaluate the performance of omicsGAT Classifier on cancer outcome prediction. There are 331 Estrogen Receptor positive (ER+) and 80 ER negative (ER-) samples, 65 Triple-negative (TN) and 346 non-TN samples in the dataset. The two tasks were to predict the ER and TN statuses of the breast cancer patients. omicsGAT Classifier was compared with SVM, RF, DNN, and GCN. First, the dataset was divided into pre-train and test set containing 80% and 20% of the total samples respectively. Then the pre-train set was divided into training and validation set containing 80% and 20% samples of the pre-train set respectively. The hyper-parameters of the proposed model used in these two tasks are listed in Supplementary Table S1. They were selected through grid search on the validation set. The same validation set was also applied to select the best model for DNN and GCN. We ran omicsGAT Classifier and baseline methods with above mentioned dataset splitting 50 times. The average AUROC scores for both omicsGAT and baseline methods are reported in Table 2. As we can see, our proposed model out-performs all the baselines for both ER and TN status predictions. Moreover, the gain in AUROC caused by omicsGAT is significant for all baselines, except SVM and RF for ER prediction. omicsGAT Classifier also offers a lower standard deviation which signifies a more consistent and stable prediction compared to the baselines. The stability of our proposed model can be pertained to the use of several heads which can secure information from different directions and the model can effectively combine them by learning distinct attention parameters for each head.

**Table 2.** The classification performance on TCGA breast cancer (BRCA) dataset. The mean AUROC scores and standard deviation (SD) of classifying patients in breast cancer subtypes are reported. *Denotes the difference between the results of omicsGAT and baseline method to be statistically significant (*p-value <* 0.001)

| Cancer Subtype | Method | AUC score | SD |
| --- | --- | --- | --- |
| ER | SVM | 0.9155 | 0.4868 |
|  | Random Forest | 0.9206 | 0.0436 |
|  | DNN | 0.8705* | 0.0586 |
|  | GCN | 0.8835* | 0.0612 |
|  | **omicsGAT** | **0.9407** | **0.0360** |
| TN | SVM | 0.8800* | 0.0722 |
|  | Random Forest | 0.8567* | 0.0663 |
|  | DNN | 0.8226* | 0.0819 |
|  | GCN | 0.8560* | 0.1023 |
|  | **omicsGAT** | **0.9368** | **0.0375** |

To evaluate the performance of omicsGAT in greater depth, the patient’s overall survival time and disease-free time were predicted on the breast cancer dataset. The Cox proportional hazards model with elastic net penalty (35) evaluated the correlation between the patient’s overall survival time or disease-free time and genomic features, i.e., the original gene expression and the omicsGAT learned embeddings. 80% of the patient samples were applied to train the model and the performance was tested on 20% test samples. The low and high risk groups on the independent test set were generated based on the prognostic index (36). The survival and disease-free prediction were visualized by Kaplan-Meier plots and compared by the log-rank test. The Kaplan-Meier plots in Figure 2 illustrates the improved patient survival time and disease-free time prediction on breast cancer patients using the embeddings generated by omicsGAT compared to the original gene expression. The log-rank test *p*-values clearly demonstrate a strong additional prediction power of the learned embeddings beyond the gene expression.

**Fig. 2.**
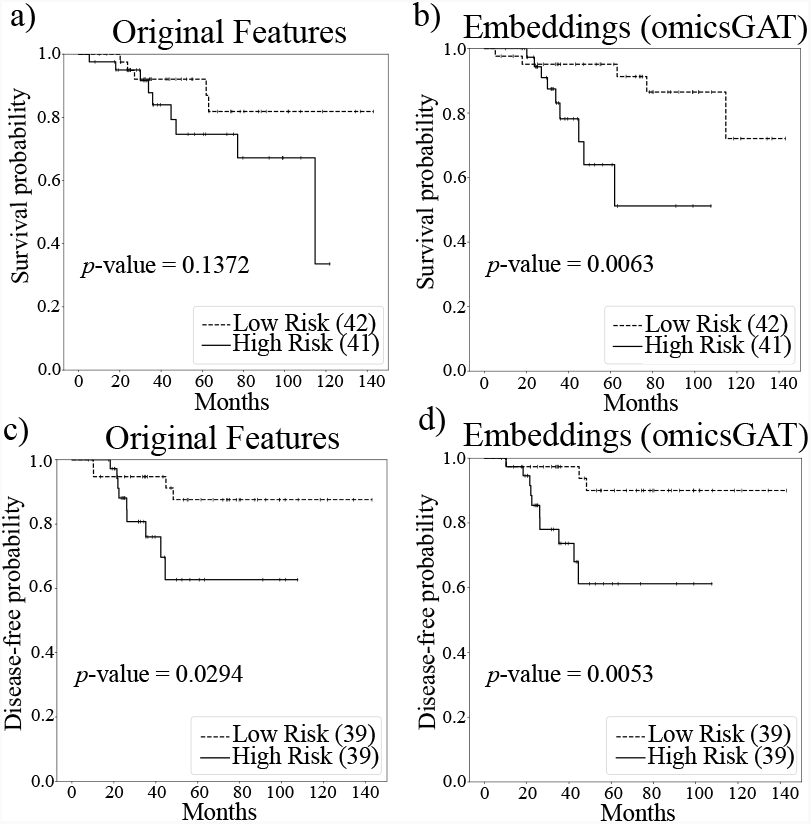
Survival and disease-free time predictions on breast cancer patients with original gene expression and the embeddings generated by omicsGAT. Kaplan-Meier plots for low (dashed line) and high (solid line) risk groups generated by a) original gene expression and b) omicsGAT learned embeddings for survival analysis; c) original gene expression and d) omicsGAT learned embeddings for disease-free analysis. The number in the parenthesis indicates the number of samples in low or high risk group. The *p*-value is calculated by the log-rank test to compare the overall survival or disease-free probability of two groups of breast cancer patients.

#### F.3. omicsGAT Improved Cancer Patient Stratification

To evaluate the generalization of our embedding mechanism, we employed omicsGAT Clustering to stratify bladder cancer (BLCA) patients. The dataset consisted of five cancer subtypes and our task was to cluster the patients into these five categories. Embeddings were generated following the first step of omicsGAT Clustering, i.e., the autoencoder described in Section C. First, the dimension of the raw gene expression data was reduced using PCA implemented through *sklearn.decomposition.PCA* package. The top 400 PCA components were then used as input in the omicsGAT pipeline, and the generated embeddings were fed to the second step of omicsGAT Clustering, a hierarchical clustering model. We show two findings in this experiment: (1) clustering of the embeddings demonstrates cluster-specific patterns in the embeddings, and (2) high-quality embeddings enhance the performance of clustering cancer patients into cancer subtypes. The embedding clustering result is illustrated in Figure 3 where each row represents a patient sample and columns represent embeddings. The patient samples were grouped together according to their cancer subtypes. The distinct pattern can be observed for the embeddings generated for a particular cancer subtype signifying the ability of omicsGAT to effectively integrate neighborhood information into the embedding for a better predictive signature.

**Fig. 3.**
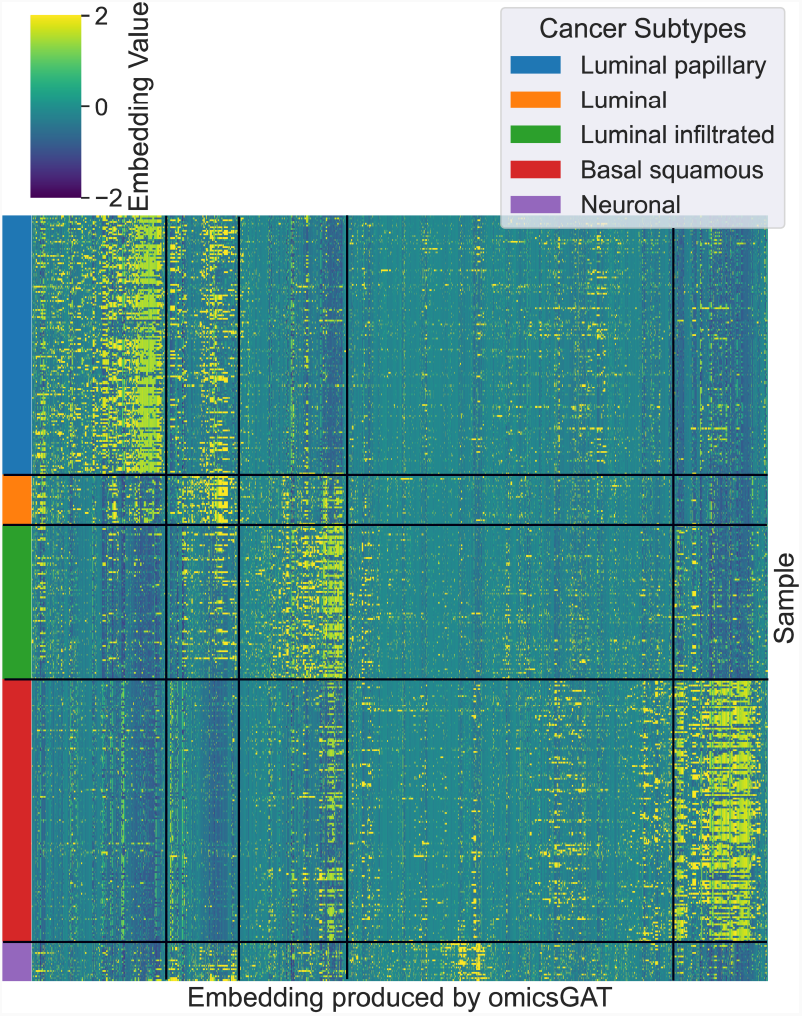
Embeddings generated by omicsGAT clustered into the corresponding cancer subtypes

Next, we compared the performance of omicsGAT Clustering with the baselines for clustering patient samples into cancer subtypes. Assigned cluster values of the samples by omicsGAT Clustering and the true cancer subtype of the samples were matched to calculate NMI and ARI scores. The NMI and ARI scores which were calculated after employing hierarchical clustering and k-means clustering on raw gene expressions, and the 400 PCA components were used as baselines. Additionally, we clustered the sample adjacency matrix

***A*** as another baseline. The results are reported in Table 4. It can be observed that both NMI and ARI scores are highest for omicsGAT Clustering followed by the clustering of the adjacency matrix. The scores for the PCA components and the raw gene expression features are lower which can be attributed to the absence of sample similarity information in the datasets, whereas the embeddings from omicsGAT and the adjacency matrix consider the relations between samples. omicsGAT used the information from the neighbors more effectively by assigning different attention coefficients to the neighbors of a sample, thereby capturing the hidden relations between samples in the embeddings. This influx of information caused by the attention mechanism in embedding generation enabled omicsGAT Clustering to outperform all baselines by a considerable margin.

**Table 3.** Hyperparameter selection for omicsGAT Clustering

| Hyperparameter | Selection Set |
| --- | --- |
| No. of PCA components (features) selected | [50, 100, 200, **400**] |
| Embedding size of a head | [4, 8, 16, 32, **64**] |
| No. of heads | [4, 8, 16, 32, **64**] |
| Network density of adjacency matrix | [0.02, 0.04, 0.1, **0.2**] |
| No. of FC layers | [2, **3**, 4] |

**Table 4.** The clustering performance on TCGA bladder cancer (BLCA) dataset. The NMI and ARI scores of omicsGAT Clustering and baseline methods are reported in the table. Hierarchical clustering was computed with ‘Manhattan’ distance and ‘Average’ linkage. Mean NMI and ARI scores with standard deviation are reported for k-means clustering (run 10 times).

| Input Data (Clustering Method) | NMI | NMI SD | ARI | ARI SD |
| --- | --- | --- | --- | --- |
| gene expression (hierarchical) | 0.0515 | - | 0.0153 | - |
| gene expression (k-means) | 0.4944 | 0.0171 | 0.4468 | 0.0548 |
| PCA components (hierarchical) | 0.1222 | - | 0.0353 | - |
| PCA components (k-means) | 0.4883 | 0.0176 | 0.4338 | 0.0388 |
| adjacency matrix (hierarchical) | 0.5448 | - | 0.5505 | - |
| **omicsGAT embeddings (hierarchical)** | **0.6147** | - | **0.6698** | - |

To visualize the clustering performance, tSNE plots (Python *seaborn* package) were created on the PCA components and the embeddings generated by omicsGAT in Figure 4 (a) and (b) respectively. Figure 4 (a) illustrates that PCA components cannot properly separate the five clusters. Although there is some separation among the patient samples in ‘Basal squamous’, ‘Luminal Papillary’, and ‘Luminal infiltrated’ subtypes, the samples in ‘Luminal’ and ‘Neuronal’ subtypes were randomly scattered on the plot. On the other hand, Figure 4 (b) shows that omicsGAT Clustering can effectively separate all five clusters, revealing the meaningful neighborhood information contained within the embeddings. Moreover, ‘Luminal’ and ‘Neuronal’ are the subtypes with the smallest number of samples which means our proposed method particularly excels at clustering rare subtypes.

**Fig. 4.**
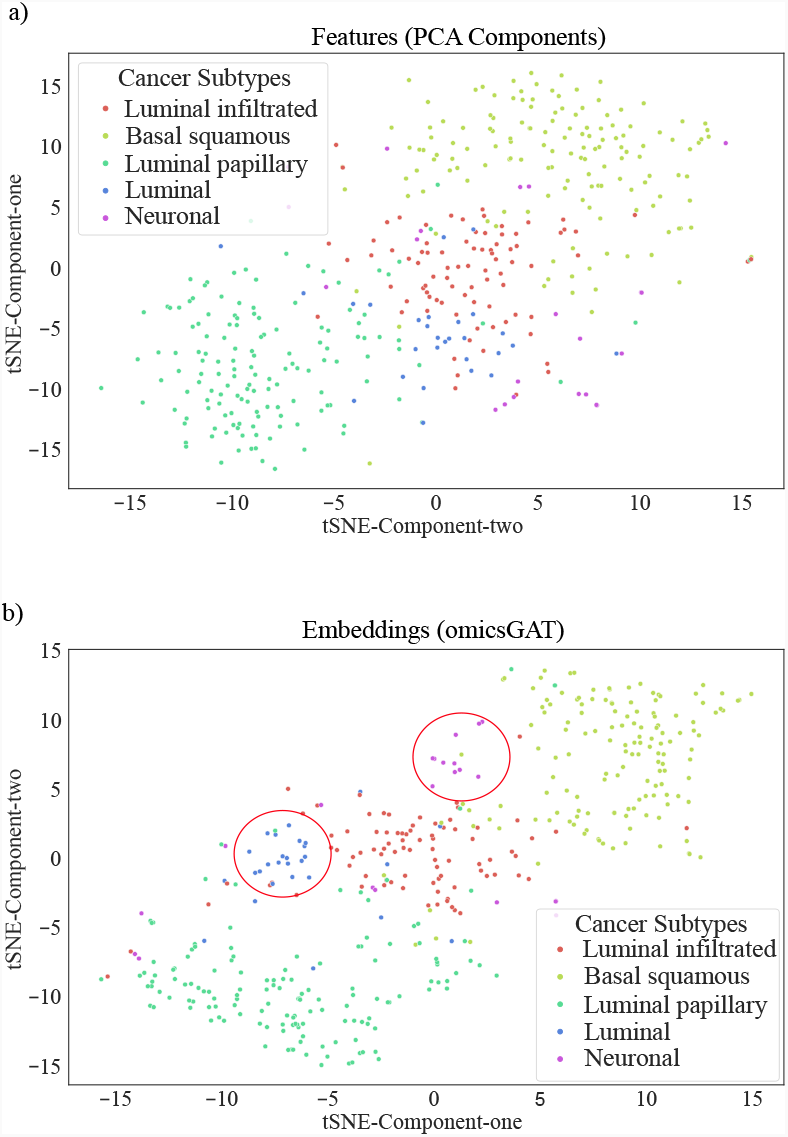
tSNE plots of the (a) PCA components generated from the BLCA data and (b) omicsGAT generated embeddings for bladder cancer patients stratification.

### G. Experimentation on Single-cell RNA-seq data

Single-cell RNA-seq (scRNA-seq) reveals heterogeneity at the cell level and offers a larger number of samples (i.e., cells) compared to bulk RNA-seq data (e.g., number of patient samples). We applied omicsGAT Clustering on scRNA-seq data and clustered cells to evaluate the generalization of our proposed model.

#### G.1. Dataset and Preprocessing

scRNA-seq data from six primary H3K27M-gliomas (H3 lysine27-to-methionine mutations) was used in the following experiment. This type of gliomas (malignant tumors) primarily arise in the midline of the central nervous system of young children (37). Early detection of tumors may improve disease prognosis; hence, stratifying the tumor cells into the correct gliomas could be very helpful for clinicians. Gene expression and label information of 2,458 cells was used for this experiment. The dataset was downloaded from the Single Cell Portal (38) and the cells were generated from six different gliomas: BCH836, BCH869, BCH1126, MUV1, MUV5, MUV10. *log*2(*x* + 1) transformed TPM (Transcripts-per-million) value was used in the analysis.

#### G.2. Single Cell Clustering

The same omicsGAT Clustering method described in Section C is followed to cluster the cells with scRNA-seq data. The top 200 PCA components were selected as the input of the omicsGAT Clustering to generate embeddings. The omicsGAT’s hyperparameters for this experiment are listed in Table S2 in the Supplementary document. The autoencoder was trained following the same steps as explained in Section F.3. Embeddings generated from the autoencoder were then fed into the hierarchical clustering model. Hierarchical and k-means clustering methods on raw gene expression and 200 PCA components were considered as the baselines along with hierarchical clustering on the adjacency matrix. As reported on Table 5, omicsGAT Clustering outperforms all the baselines which means the cluster assignments resulting from the omicsGAT generated embeddings are more similar to the true label information. This result is corroborated by the tSNE plots in Figure 5 (a) and (b) which are drawn on the PCA components and the embeddings generated by omicsGAT respectively. The tSNE plot for omicsGAT Clustering shows more separation among the clusters as compared to the PCA components. Specifically, for the ‘MUV1’ group, our model formed a single cluster containing all the cells belonging to that type (red circle in Figure 5 (b)), whereas the tSNE plot using PCA components shows two different clusters for the cells in ‘MUV1’. Based on the results, we can conclude that in the case of scRNA-seq data analysis, omicsGAT Clustering can take advantage of the detailed cellular level information and uses the attention mechanism on the cell-cell similarity network to better cluster the samples.

**Table 5.**
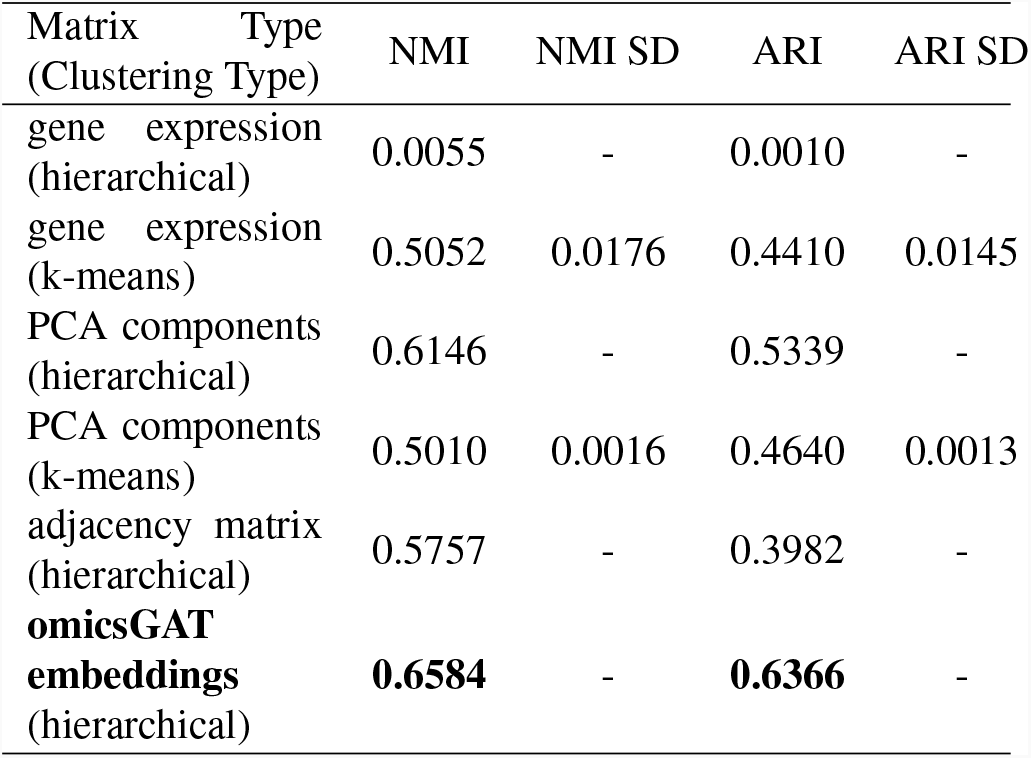
The clustering performance on scRNA-seq H3K27M-gliomas data. The NMI and ARI scores of omicsGAT Clustering and baseline methods are reported in the table. Hierarchical clustering was computed with ‘Cosine’ distance and ‘Average’ linkage. Mean NMI and ARI scores with standard deviation are reported for k-means clustering (run 10 times).

**Fig. 5.**
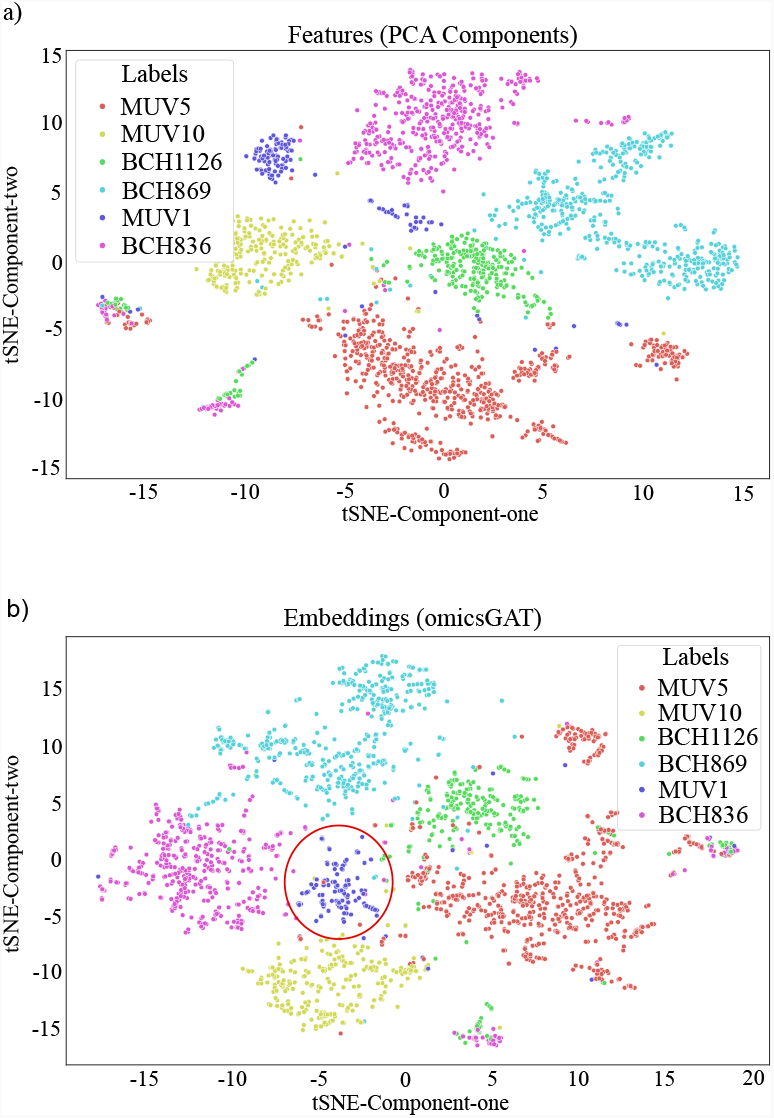
tSNE plots of the (a) PCA components generated from the scRNA-seq data and (b) omicsGAT generated embeddings for cell clustering.

## Discussion

omicsGAT can successfully integrate the structural information within gene expression data into sample embeddings enabling better classification and clustering performance compared to the original dataset. The stronger predictive ability of the embeddings is contributed by the self-attention mechanism in omicsGAT. A binary adjacency matrix is applied to define neighborhoods in omicsGAT that includes self-connections to ensure that the information of a sample itself is also considered in the embedding. The performance is reduced when we ran the same classification task with just the adjacency matrix. The adjacency matrix is calculated using correlation only, which keeps track of the pairwise linear relations between samples. However, using the attention mechanism, omicsGAT can capture complex nonlinear relations by accounting for the importance of neighboring samples on the classification or clustering of a target sample. The captured relations among samples are represented in the generated embeddings which enables the model to perform better on classification and clustering tasks.

In order to verify the effect of the multi-head attention mechanism, a *sample* × *sample* attention matrix was constructed by extracting the attention coefficients from a trained omicsGAT model following the method used by Ullah and Ben-Hur (39). For a target sample, each of the *h* heads assigns different attention coefficients to its neighbors, and only the highest among the *h* attention coefficients was considered for each neighbor to represent its relation with the target sample. The same procedure is repeated to generate the full attention matrix. This process was applied to build the attention matrix for both BLCA and cell clustering tasks described in Section F.3 and Section G.2 respectively. This attention matrix reveals the importance of combining the attention mechanism with the network information received through the adjacency matrix. As seen in Table 6, clustering on the attention matrix outperforms the clustering on the adjacency matrix for both datasets. Moreover, the clustermap of the attention matrix obtained from the trained model on BLCA data, illustrated in Figure 6, shows a distinct pattern of the cancer subtypes specifically for ‘Luminal papillary’ and ‘Basal squamous’. From these results, we can conclude that some neighbors play a more important role than others in classification or clustering of a sample, and omicsGAT can effectively inject this information into the model along with the graph structure to generate more meaningful embeddings for better downstream analyses. An important aspect of omicsGAT is the use of multiple heads. The learnable weight parameters (***W*** and ***a***) of each head were initialized separately using the *xavier normal* library function in *Pytorch* (31).

**Table 6.** NMI and ARI scores of the Hierarchical Clustering applied on attention and adjacency matrices

| Dataset | Input Matrix | NMI | ARI |
| --- | --- | --- | --- |
| BLCA | adjacency matrix | 0.5448 | 0.5505 |
|  | attention matrix | 0.5743 | 0.6373 |
| scRNA | adjacency matrix | 0.5757 | 0.3982 |
|  | attention matrix | 0.5788 | 0.4821 |

**Fig. 6.**
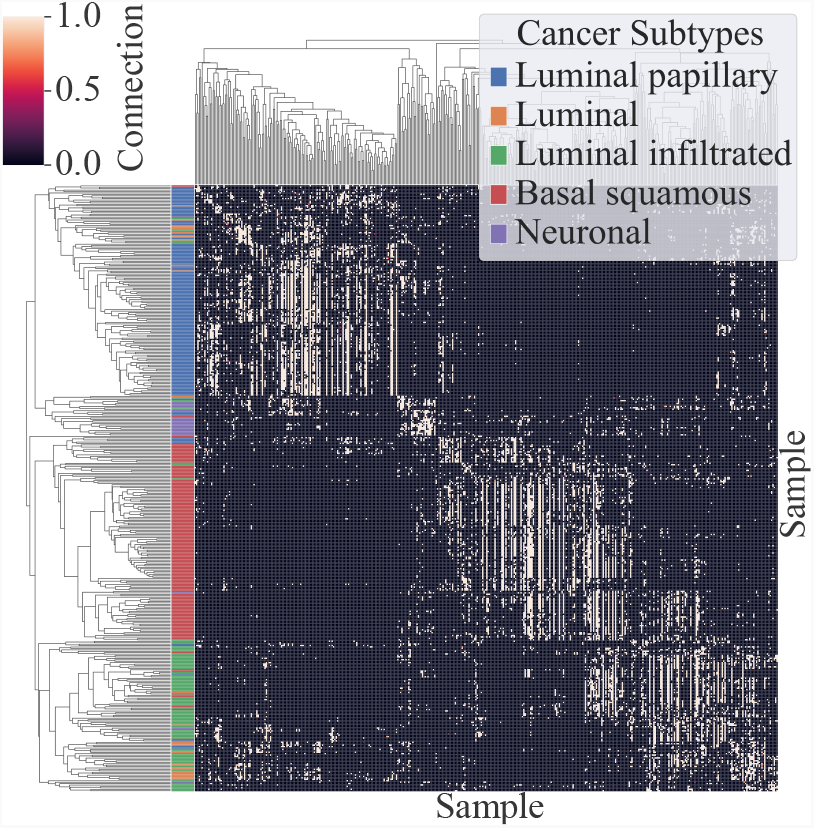
Clustermap of the Attention Matrix generated from the trained omicsGAT model on BLCA data

For the clustering tasks, the NMI and ARI scores of the baselines were relatively low with hierarchical clustering which can be observed in Table 4 and Table 5. Therefore, we also applied k-means clustering to them in order to compare them with omicsGAT. Since the performance of k-means clustering depends on the initialization of the cluster-centers, clustering was conducted 10 times and the mean scores along with standard deviations were reported in the tables.

## Conclusion

Powered by high-throughput genomic technologies, the RNA-seq method is capable of measuring transcriptome-wide mRNA expressions and molecular activities in cancer cells. Hundreds of computational methods have been developed for cancer outcome prediction, patient stratification, and cancer cell clustering. Some of these methods consider sample-sample similarities in the analysis, and some of them do not. These sample similarity-based methods cannot distinguish the importance of the neighbors for a particular sample in the downstream prediction or clustering tasks. Therefore, we introduced omicsGAT in this study which leverages a self-attention mechanism consisting of multiple heads to assign proper attention weights to the neighbors of a sample in the network. Experiments on cancer subtype analyses show the superior performance of the model in every aspect compared to the baseline methods. We also show the generalization of omicsGAT’s performance on both bulk RNA-seq and scRNA-seq data. As a future objective, we would like to extend omicsGAT to include metapath selection which would consider the best paths in a network to perform a certain task on a particular sample.

