## Supplementary document for "omicsGAT: Graph Attention Network for Cancer Subtype Analyses"

### 1 Tables

Table S1: Sets of hyperparameters in omicsGAT for cancer outcome prediction

| Hyperparameter | Selection Set |
|---|---|
| No. of features selected using correlation | $[128, 256, 512, 1024]$ |
| Embedding size of a head | $[4, 8, 16, 32, 64]$ |
| No. of heads | $[4, 8, 16, 32, 64]$ |
| No. of neighbors of a sample in the adjacency matrix | $[5, 10, 20, 40]$ |

Table S2: Hyperparameter selection for single-cell clustering. The bold hyperparameters are selected for the scRNA-seq clustering task based on the least reconstruction loss of the decoder.

| Hyperparameter | Selection Set |
|---|---|
| No. of PCA components (features) selected | $[50, 100, \mathbf{200}, 400]$ |
| Embedding size of a head | $[4, \mathbf{8}, 16, 32, 64]$ |
| No. of heads | $[4, \mathbf{8}, 16, 32, 64]$ |
| No. of neighbors of a node in adjacency matrix | $[5, \mathbf{10}, 20, 40]$ |
| No. of FC layers | $[2, \mathbf{3}, 4]$ |
